# Neutrophil Peptidylarginine Deiminase 4 is Essential for Detrimental Age-related Cardiac Remodeling & Dysfunction in Mice

**DOI:** 10.1101/2022.03.14.484062

**Authors:** Stijn Van Bruggen, Sirima Kraisin, Jore Van Wauwe, Paolo Carai, Liesbeth Frederix, Thilo Witsch, Kimberly Martinod

## Abstract

**Aims:** We aimed to study the long-term effect of neutrophils on cardiac health during the process of natural aging. We hypothesized that neutrophil PAD4, via its role in neutrophil extracellular trap (NET) formation, is involved in myocardial remodeling and cardiac fibrosis development, resulting in turn in impaired cardiac function.

**Methods and results:** We generated mice with deletion of *Padi4*, a NET-essential gene, under the neutrophil-specific promoter S100A8 (PAD4^fl/fl^MRP8Cre^+^). These mice and their littermate controls were aged for two years (coinciding with approximately 70 years of age in humans; the age at which HF is the number one cause of hospitalization), after which cardiac function and remodeling were evaluated. We performed a comprehensive echocardiography analysis including both structural and functional parameter measurements. Deletion of PAD4 in neutrophils resulted in a protection against both systolic, and diastolic dysfunction. Interestingly, these mice showed protection against age induced fibrosis, detected as through the absence of cardiac collagen deposition. To explore this further, cardiac gene expression and plasma cytokine levels were evaluated. Here we saw a clear impact of PAD4-deficiency on cardiac neutrophil recruitment, with both cardiac genes as well as plasma cytokines involved in neutrophil recruitment being downregulated in aged PAD4^fl/fl^MRP8Cre^+^ animals in comparison to littermate PAD4^fl/fl^ controls, including decreased plasma levels of C-X-C ligand 1 (CXCL1).

**Conclusion:** Our data confirms neutrophil PAD4 involvement in heart failure progression by promoting cardiac remodeling, leading to cardiac dysfunction with old age. We saw that the deletion of PAD4 specifically in neutrophils had an influence on the CXCL1-CXCR2 axis, which is known to be involved in HF development.

**Translational perspective:** In the developed world, an estimated 2% of the population lives with heart failure (HF). HF can be viewed as an upcoming pandemic, which is only expected to increase due to the aging of the global population. Therefore, research in HF development and progression are needed to establish new avenues for treatment and improved therapies. In our study, we were able to show the contribution of neutrophil PAD4 to HF pathogenesis, providing new supporting evidence for the involvement of NETs in detrimental cardiac remodeling.

## Introduction

According to projections from the World Health Organization (WHO), by 2030 one in six people will be 60 years or older. By 2050, this will have increased to one in five. This will undoubtedly have a profound societal impact and underscores the need for research into healthy aging. Aging can be seen as a complex interplay between environmental, stochastic, genetic, and epigenetic factors. Increasing age is often accompanied by a chronic low-grade pro-inflammatory status, even in the absence of any form of infection. This state of sterile inflammation with increasing age has been defined as “inflammaging” ^1^. Chronic inflammation, as evaluated by plasma or serum levels of pro-inflammatory mediators, can cause malfunctioning of several cellular and molecular events, ultimately leading to various chronic ailments and diseases, as well as the loss of tissue integrity and organ function over time ^2-4^. This organ damage can result in age-related pathologies, e.g. Alzheimer’s disease, atherosclerosis, arthritis, cancer, and cardiovascular diseases ^5, 6^.

A final pathological outcome of many of these diseases is the development of fibrosis, also known as pathological collagen deposition ^7^. Although the deposition of collagen, and other extracellular matrix (ECM) proteins is essential in wound healing, normal tissue repair can evolve into an uncontrolled, irreversible fibrotic response if tissue injury is either too severe, or repetitive ^8^. During fibrosis development, also known as fibroplasia, connective tissue replaces normal parenchymal tissue ^8^. Even though this process initially starts off being beneficial for organ function and healing, during chronic inflammation the repair process becomes inappropriately controlled and pathogenic, resulting in substantial ECM production and deposition, ultimately leading to the replacement of normal tissue by a fibrotic scar ^9^.

Fibrosis development is an extremely complex and multi-stage process, in which bone marrow-derived leukocytes play an essential role ^10^. Early on during tissue injury, platelets degranulate and release chemotactic molecules and growth factors, including transforming growth factor beta-1 (TGF-β1) and platelet derived growth factor (PDGF) ^11^. Additionally, activated platelets which have externalized their granules have higher expression of P-selectin on their surface, which is capable to interact with P-selectin granulocyte ligand 1 (PSGL1), constitutively expressed by neutrophils and other leukocytes ^12, 13^. This interaction results in platelet-neutrophil complex formation, followed by neutrophil activation ^14^. As a result of this platelet-leukocyte interaction, the neutrophil metabolic state is altered, resulting in the release of granule proteins ^15^, enhanced phagocytic capabilities ^16^, and the production and release of reactive oxygen species (ROS) ^17^.

In neutrophils, ROS production can result in the disruption of azurophilic or primary granules, causing the cytoplasmatic release of proteases like myeloperoxidase (MPO) and neutrophil elastase (NE) ^18^. These proteins can migrate to the nucleus, with MPO facilitating the initial entry of NE past the nuclear membrane. In the nucleus, NE starts degrading histones, promoting chromatin decondensation and nuclear swelling ^19^. This increase in nuclear volume continues until both nuclear, and plasma membranes are incapable of resisting tensile stress, leading to cell rupture. The loss of membrane integrity is coupled with the release of nuclear material, lined with a range of proteins, including MPO, NE, and histones. These extracellular DNA structures are known as neutrophil extracellular traps (NETs) ^20^. This process of NET formation (also known as NETosis), requires the activation of peptidylarginine deiminase 4 (PAD4). Equipped with a nuclear localization sequence (NLS), PAD4 acts by citrullinating arginine residues on proteins, resulting in the overall loss of positive charges of these amino acid residues ^21^. When reaching the nucleus, PAD4 can citrullinate specific arginine residues on histone tails, facilitating chromatin decondensation, a process essential during NETosis ^21^. Although NETs were first described as an anti-bacterial defense mechanism ^22, 23^, NETosis can also occur during sterile inflammation, as is the case during aging ^24^. Once released, NETs can cause damage to underlying tissue, and are both proinflammatory and prothrombotic ^25-27^. Furthermore, NETs are released in a range of pathological conditions, including deep vein thrombosis ^25, 27^, cancer ^26, 28^, myocardial ischemia/reperfusion injury ^29^, atherosclerosis ^30-32^, rheumatoid arthritis ^33^, and other auto-immune diseases ^34^. In all these pathologies, NETs have been established as having a direct contribution to pathophysiology, and/or a detrimental effect on disease progression. Additionally, it was shown that NETs are capable of catalyzing the conversion of fibroblasts towards collagen secreting myofibroblasts *in vitro* ^29^, thus directly linking NETs to fibrosis development. However, this has not yet been demonstrated in the heart.

We have previously studied the interplay between NET release, aging, and cardiac fibrosis by studying the effect of systemic deletion of PAD4 in aged mice ^24^. This resulted in a cardioprotective phenotype with increasing age, with preservation of both systolic, and diastolic function. It was shown in the same study that, with PAD4 deletion, excessive deposition of interstitial cardiac fibrosis was absent with increasing age. However, the phenotype could not be specifically attributed to neutrophils and NETs. Therefore, the overall goal of this study was to selectively knock out PAD4 in neutrophils, and to investigate how neutrophil PAD4 is mechanistically involved in the complex process of spontaneous fibrosis development with increasing age.

## Materials and Methods

### Animals

B6.Cg-Padi4^tm1.2Kmow^ (PAD4fl/fl, RRID **IMSR_JAX:026708**) mice were purchased from the Jackson Laboratory (USA) and backcrossed for seven generations with C57BL/6J mice purchased from Charles River (France). Intercrossing of these mice with B6.Cg-Tg(S100A8-cre,-EGFP)1Ilw/J (MRP8-Cre-ires/GFP, RRID **IMSR_JAX:021614**) obtained from the Jackson Laboratory (USA) resulted in PAD4fl/fl x MRP8-Cre-ires/GFP (PAD4fl/flMRP8Cre+). Breeding of PAD4fl/fl with PAD4fl/flMRP8Cre+ mice resulted in litters containing both PAD4^fl/fl^ and PAD4^fl/fl^MRP8Cre^+^ offspring due to the MRP8Cre hemizygosity.

Knockout mice (PAD4^fl/fl^MRP8Cre^+^), together with littermate controls (PAD4^fl/fl^) were aged for 24 months (mo). Separate groups of 9 to 12 week-old mice from the same breeding colony were used as controls. Mice were kept on a standard laboratory diet (ssniff #R/M-H) for the entirety of the study. All groups were age- and sex-matched and received *ad libitum* feed, with free access to water. All experimental procedures were reviewed and approved by the Ethical Committee of the Laboratory Animal Center at the KU Leuven (Project number P019/2020), according to the Belgian Law and the guidelines from Directive 2010/63/EU of the European Parliament. All animal interventions were performed during morning hours in order to take circadian rhythms of both mice and neutrophils into account.

### Obesity and hypertension-induced HFpEF

At two years of age, mice were changed from regular chow to high fat diet feed containing 60 kcal% fat (Research Diets #D12492). Additionally, N^ω^-nitro-L-arginine methyl ester (L-NAME) (Sigma-Aldrich) was added to the drinking water (0.5 g/L) ^35^. These metabolic stressors were administrated to induce further diastolic dysfunction. Mice were monitored daily, and kept on the diet for six weeks. Cardiac function was assessed every three weeks, starting the day of diet administration.

### Echocardiography

Cardiac function and dimensions were measured via echocardiography, using a Vevo 2100 3D analyzer (Fujifilm Visualsonic). Mice were anesthetized using 2% isoflurane in medical oxygen at a flow rate of 2.5 L/min. Body temperature was constantly monitored via an anal probe and kept between 35.5°C and 37°C. Heart rate was kept stable between 450 and 550 BPM for all measurement acquisitions. In the Parasternal Long Axis (PLAX), cardiac dimensions were evaluated using Brightness (B)-mode. Using Pulsed wave (PW) doppler the blood flow, and pressure in the pulmonary artery was measured. In the parasternal short axis (PSAX) Motion (M)-mode was used to measure left ventricular posterior wall (LVPW) thickness, left ventricular internal diameter (LVID), and left ventricle anterior wall (LVAW) thickness. In the apical 4 chamber (A4C) window, PW doppler was used at the height of the mitral valve to measure blood flow in the left ventricle, coming from the left atrium. Using PW Doppler in the pulmonary artery, right ventricular pulmonary ejection time (PET) and pulmonary acceleration time (PAT) were measured. Finally, blood flow in the aorta was measured using PW doppler. Echocardiographic recordings were stored digitally, and analyzed using the Vevo Lab software (Vevo lab, V5.5.1). Left ventricular ejection fraction (LVEF) was calculated based on Simpson’s method. Blood flow in the A4C view was used to determine signs of impaired LV relaxation, as ease of ventricular filling is expressed as the ratio between the E and the A wave (E/A).

### Plasma preparation

At time of euthanasia, mice were anesthetized using a mixture of ketamine/xylazine (125 mg/kg and 12.5 mg/kg) at a non-lethal dose. Once mice were non-responsive to pedal-reflex (toe pinch), blood was collected retro-orbitally in 3.8% citrate anti-coagulant in a 1/10 dilution using a pre-coated capillary. After collection, blood was centrifuged at 3000 g for five minutes after which the supernatant was transferred to a clean tube and centrifuged again for five minutes at 12300 g. After centrifugation, platelet-poor plasma was transferred to a clean tube and immediately stored at -20°C for future batch analysis.

### Histology

Ketamine/xylazine anesthetized mice were perfused using 0.9% saline until liver paleness was verified, after which organs were removed. Organs were fixed in 4% paraformaldehyde in phosphate-buffered saline (PBS) overnight at 4°C. After washing 3 times in PBS, fixed organs were kept in 70% ethanol at 4°C until further processing and paraffin embedding. Tissue was sectioned in 8 µm slices and rehydrated. To assess collagen content in heart tissue, Masson Trichrome (Sigma Aldrich) and Fast Green/Sirius Red (Chondrex) staining were performed according to the manufacturer’s protocols. After staining, slides were dehydrated and mounted using DPX mounting medium (sigma). Heart sections were then visualized at 100X magnification in bright-field microscopy in a blinded manner. MosaicX images were acquired using Axiovision software. These images were used to quantify collagen content as a percentage of the total area using color thresholding analysis in ImageJ software (FIJI) ^36^.

### RT-qPCR

Heart tissue was snap frozen in liquid nitrogen, and subsequently stored at -80°C. Later, tissue was mechanically grinded and homogenized using ceramic beads. Homogenized tissue was used for total RNA extraction via an RNeasy mini kit (QIAGEN). Total RNA was used to construct cDNA using random hexamer primers. For cDNA construction, QuantiTect Reverse Transcription kit, with gDNA removal step (QIAGEN) was used according to the manufacturer’s instructions. Quantitative real-time PCR was performed using gene specific primes (supplemental table 1). SYBR Green Master Mix (Applied Biosystems) was used to perform qRT-PCR with a QuantStudio™ 3 Real Time PCR detection system. RT-qPCR data was analyzed using the Livak and Schmittgen (2^-ΔΔCt^) method ^37^.

### Plasma analyses for biomarker determination

Stored plasma was used for detection of several inflammation biomarkers. MPO-DNA ELISA was performed as previously described ^38^. Plasma levels of cell free dsDNA were detected using the Quanti-iT™PicoGreen™ dsDNA assay kit, according to the manufacturers instructions (ThermoFisher). Mouse IL-6 levels in plasma were initially detected using the ProQuantum Immunoassay kit for detection of mouse Il-6, according to manufacturer’s instructions (ThermoFisher).

For multiplex cytokine detection, the V-PLEX Cytokine Panel 1 mouse MSD kit (Meso Scale Discovery) was run according to the manufacturers instructions. This assay allowed for high-sensitivity detection of IL-15, IL-17A/F, IL27p28/IL-30, IL-33, IP-10, MCP1, MIP1a, and MIP2. For multiplex detection of pro-inflammatory markers, V-PLEX Proinflammatory Panel 1 mouse kit (MSD) was performed according to manufacturer’s instructions. This assay allowed for the detection of IFN-γ, IL-1β, IL-2, IL-4, IL-5, IL-6, IL-10, Il12p70, KC/GRO, and TNF-α. Data of V-PLEX Meso Scale experiments were analyzed using the MSD discovery workbench software (MSD, LSR_4_0_13) according to the manufacturer’s instructions.

### Statistical analysis

Data are represented as median ± interquartile range in the figures, means ± SEM in the text. For statistical tests, a two-tailed Mann-Whitney U test was used to compare two groups. For comparison of more than two groups, Kruskal-Wallis test was applied. All tests were performed after elimination of statistical outliers, according to ROUT’s method. Correlation analysis was performed between the level of cardiac collagen, which is a measure for heart fibrosis, and both left ventricular ejection fraction (LVEF) and E/A ratio, using simple linear regression. All statistical tests were performed using Graphpad Prism software (version 9).

## Results

### Age-induced ventricular dilation is lacking in the absence of neutrophil PAD4

Aging in humans is characterized by a gradual reduction in the number of cardiomyocytes, both through necrosis and apoptosis ^39-42^. This cardiomyocyte loss results in eccentric remodeling of the left ventricle, leading to dilated cardiomyopathies (DCM) ^43^. Throughout DCM, the ventricles dilate and lose their ability to properly contract. DCM development is among the most common causes of HF development ^43^, which was recently defined and generalized by a joint effort of the European Society of Cardiology (ESC), the American Heart Association (AHA), and the Japanese Heart Failure Society ^44^. In this joint effort of standardizing heart failure definition globally, HF was defined as *“a clinical syndrome with symptoms and/or signs caused by a structural and/or functional cardiac abnormality and corroborated by elevated natriuretic peptide levels and/or objective evidence of pulmonary or systemic congestion”* ^44^.

To confirm that our model was adequate to study age-induced cardiac remodeling, ventricular dimensions of both young (9 - 12 wk old) (*PAD4*^*fl/fl*^ n = 15, *PAD4*^fl/fl^MRP8Cre^+^ n = 11) and old (24 mo old) (PAD4^fl/fl^ n = 25, *PAD4*^fl/fl^MRP8Cre^+^ n = 14) mice were evaluated. In order to study the effect of neutrophil PAD4 on this process, PAD4^fl/fl^MRP8Cre^+^ were evaluated alongside littermate controls (Fig. 1 A). Ventricular dimensions were evaluated in age- and sex-matched groups including both male and female mice. Mice were housed in the same animal room and received standard rodent chow for their entire lifespan. Using echocardiography, left ventricular dimensions were evaluated both at peak systole and diastole. During peak systole, no significant differences were observed in LV wall thickness, with both the LV anterior wall (LVAW) (Fig. 1 B), as well as the LV posterior wall (LVPW) (Fig. 1 D) thickness of the aged groups being comparable to the young controls. When looking at ventricular diameter, the LVID(s) increased significantly in the old PAD4^fl/fl^ group both compared to the young (young PAD4^fl/fl^ – 2.29 ± 0.12 mm; old PAD4^fl/fl^ – 2.88 ± 0.12 mm) control group, as well as to the PAD4^fl/fl^MRP8Cre^+^ experimental group (old PAD4^fl/fl^MRP8Cre^+^ - 2.43 ± 0.15 mm) (Fig. 1 C). During diastole, again, no change in LVAW (Fig. 1 E), or LVPW (Fig. 1 G), were detectable between young and old groups. However, the old PAD4^fl/fl^ showed a significant dilation of the left ventricle (LVID(d)) compared to the young controls (young PAD4^fl/fl^ – 3.58 ± 0.13 mm; old PAD4^fl/fl^ – 3.99 ± 0.08 mm) (Fig. 1 F). Interestingly, this dilation was absent in the old PAD4^fl/fl^MRP8Cre^+^ (old PAD4^fl/fl^MRP8Cre^+^ - 3.72 ± 0.15 mm), with internal diameter being comparable to their young controls (young PAD4^fl/fl^MRP8Cre^+^ - 3.44 ± 0.10 mm) (Fig. 1 F).

**Figure 1.**
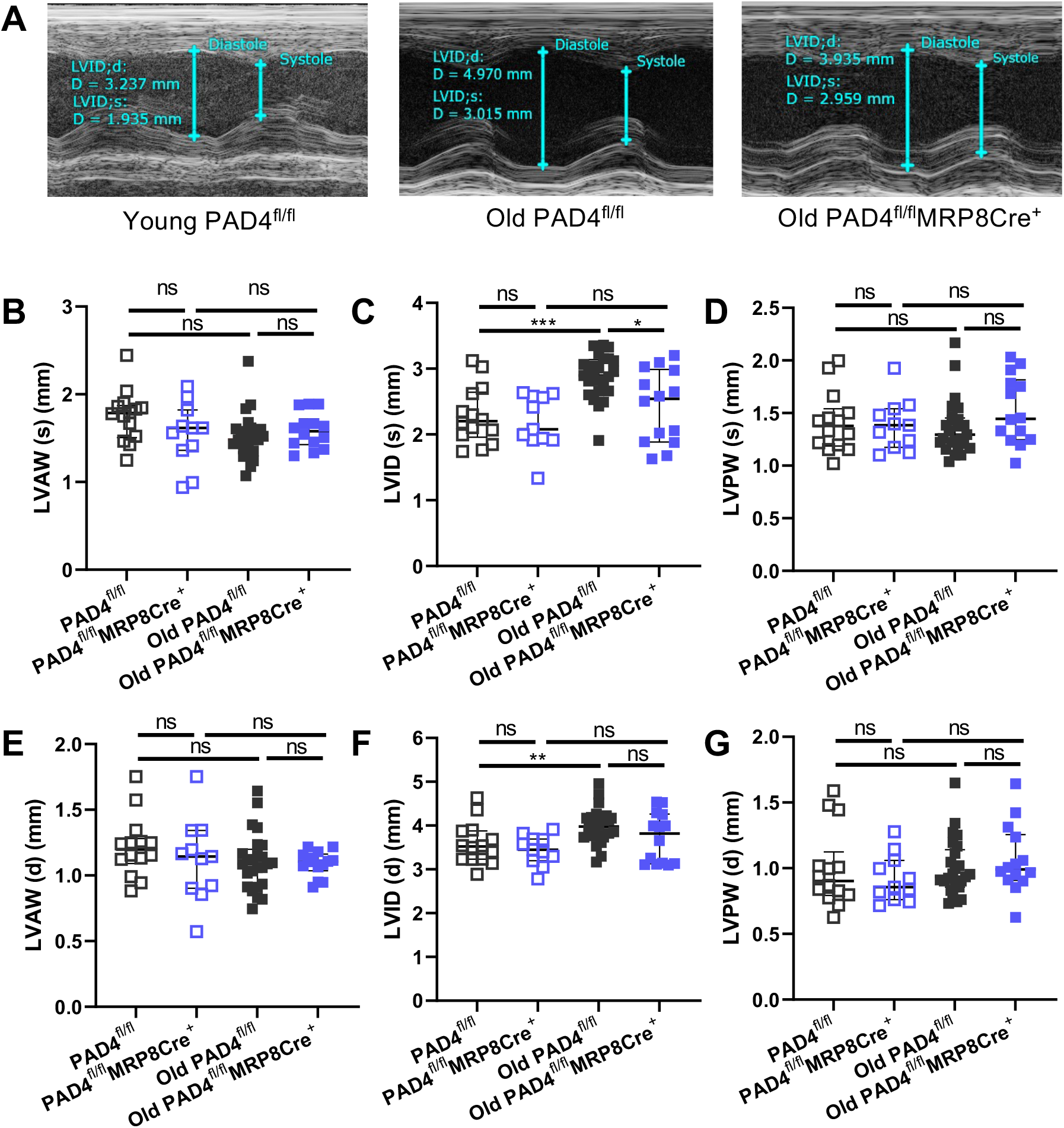
Neutrophil PAD4 deletion prevents left ventricular dilation with increasing age. (A) representative transthoracic echocardiography images of M (motion) mode recording in parasternal short axis (PSAX) window in young (9 – 12 wk) and old (24 mo) PAD4^fl/fl^ and PAD4^fl/fl^MRP8Cre^+^ mice. Dimensions are given by LV anterior wall (LVAW), left ventricular internal diameter (LVID), and left ventricular posterior wall (LVPW). (B and C) quantification of left ventricular dimensions of young and old PAD4^fl/fl^ and PAD4^fl/fl^MRP8Cre^+^ mice. NS: not significant, * P < 0.05, ** P < 0.01, *** P < 0.001. Graphs show n = 10 – 32 for young mice, and n = 14 – 25 for old mice.

### Age-induced decline in heart function is absent in PAD4^fl/fl^MRP8Cre^+^ mice

We next analyzed the effect of aging on cardiac function in both PAD4^fl/fl^ and PAD4^fl/fl^MRP8Cre^+^ mice (supplemental table 2). Systolic function was evaluated using left ventricular ejection fraction (LVEF), which is a measure for efficiency of ventricular emptying, and was estimated using Simpson’s method which considers end-systolic and end-diastolic volumes. As expected, deletion of neutrophil PAD4 did not have an impact on systolic function in young mice. In old PAD4^fl/fl^ mice, in contrast, we saw a decline in LVEF, which is consistent with results found in literature for wild-type animals ^24^. The decline in LVEF of old PAD4^fl/fl^ mice was attributed to decreased contractibility of the left ventricle, which was found to be less pronounced in old PAD4^fl/fl^MRP8Cre^+^ mice (Fig. 2 A). Interestingly, our data clearly show that LVEF in the PAD4^fl/fl^ group dropped from 67 ± 2% in the young group to 53 ± 2 % in the old PAD4^fl/fl^ group. In contrast, LVEF in the old PAD4^fl/fl^MRP8Cre^+^ group remained constant over the two-year period (young – 66 ± 2 %, Old – 67 ± 2 %) (Fig. 2 B). This is in agreement with the systemic PAD4^-/-^ model previously described ^24^.

**Figure 2.**
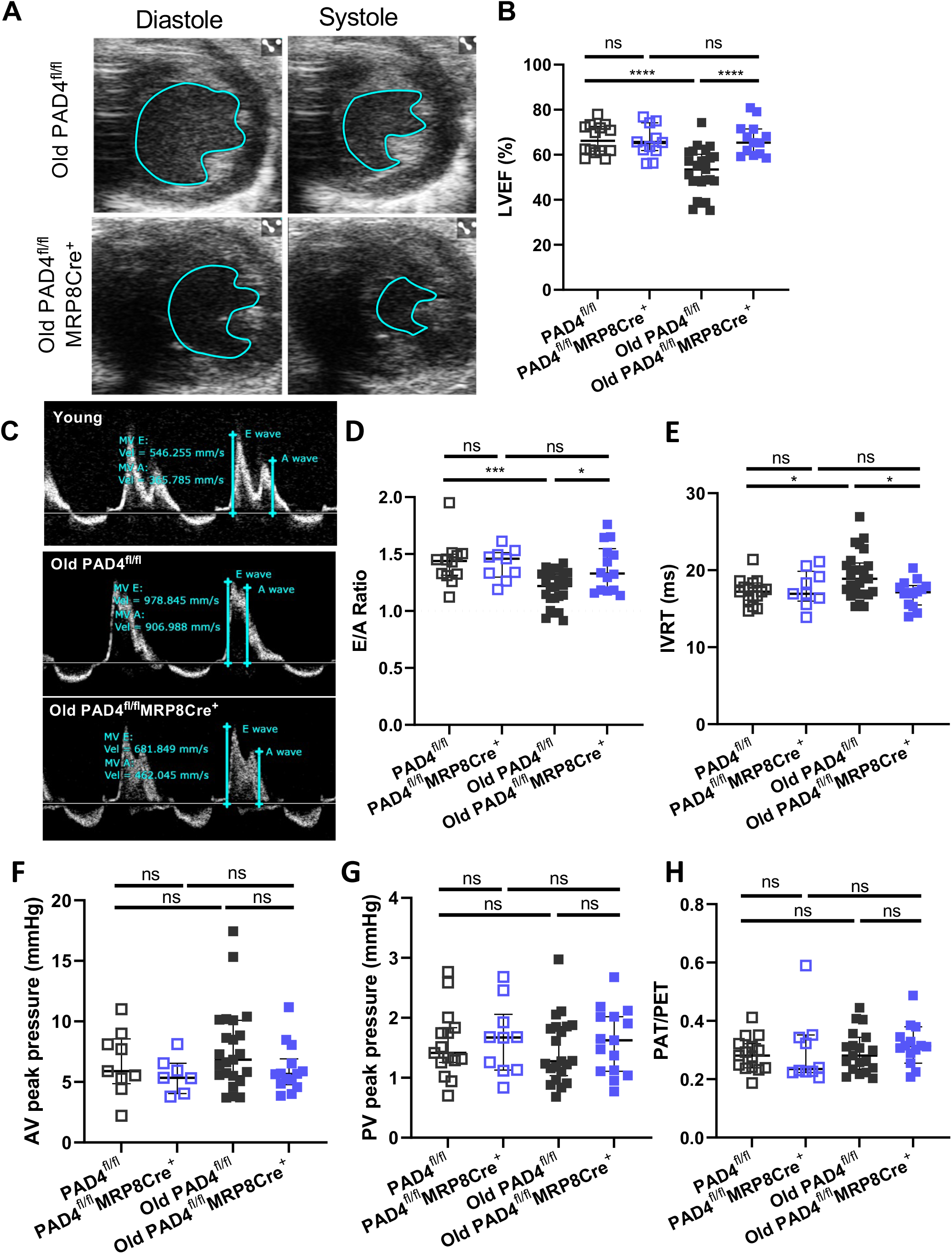
PAD4^fl/fl^MRP8Cre^+^ mice maintain cardiac function comparable to young mice. (A) Representative PSAX transthoracic echocardiography images of the left ventricle at peak diastole and systole of both old (24 mo) PAD4^fl/fl^ and PAD4^fl/fl^MRP8Cre^+^ mice. (B) Quantification of LV dimensions as a mean of showing systolic function through left ventricular ejection fraction (LVEF), measured through PSAX M-mode. (C) Representative pulsed wave (PW) Doppler echocardiography images in the apical 4 chamber (A4C) window. Ventricular diastolic function was evaluated in young and old PAD4^fl/fl^ and PAD4^fl/fl^MRP8Cre^+^ mice through the flow pattern across the mitral valve. (D and E) LV ventricular diastolic function was evaluated through the filling pattern, evaluated and calculated as the ratio between the E and A wave and the isovolumetric relaxation time as the time (IVRT) between aortic ejection and early LV filling. (D) Calculation and quantification of LV filling pattern by taking the E/A ratio. A E/A ratio equal to 1.5 (E > A) is taken as a normal pattern, while E < A is a reversed pattern. (E) Quantification of IVRT in young and old PAD4^fl/fl^ and PAD4^fl/fl^MRP8Cre^+^ mice, increasing IVRT is evidence of impaired LV filling. (F and G) quantification of peak pressure in the aorta and pulmonary artery respectively. Pressure were evaluated via PW Doppler measurements in the corresponding blood vessels through transthoracic echocardiography at peak systole. (F) quantification of peak systolic pressure at the height of the aortic valve (AV) in young and old PAD4^fl/fl^ and PAD4^fl/fl^MRP8Cre^+^ mice. (G) Quantification of peak systolic pressure at the height of the pulmonary valve (PV) in PAD4^fl/fl^ and PAD4^fl/fl^MRP8Cre^+^ mice. (H) evaluation of right heart function. Ratio between pulmonary acceleration time (PAT) and pulmonary ejection time (PET) was taken as a way of quantifying right ventricular (RV) function. NS: not significant, * P < 0.05, **** P < 0.0001. Graphs show n = 10 – 32 for young mice, and n = 14 – 25 for old mice.

Considering the effect aging has on diastolic function, atrial to ventricular filling was measured to assess diastolic function of the left ventricle. Specifically, the mitral inflow pattern was measured, and the ratio between the E-wave (representing early, passive filling of the left ventricle due to opening of the mitral valve), and the A-wave (representing late active filling due to atrial contraction) was calculated. Generally, an E/A ratio below one is considered a clinical sign for diastolic dysfunction ^45^. In the old PAD4^fl/fl^ mice, the mean E/A ratio significantly decreased compared to young PAD4^fl/fl^ mice (old PAD4^fl/fl^ 1.2 ± 0.03, young PAD4^fl/fl^ 1.5 ± 0.06) (Fig. 2 C-D), confirming the diastolic cardiac dysfunction as a result of aging. Interestingly, this evolution towards diastolic dysfunction is absent in the old PAD4^fl/fl^MRP8Cre^+^ mice, with a mean E/A ratio of 1.4 ± 0.06, corroborating the previously seen protection against cardiac dysfunction in the PAD4^fl/fl^MRP8Cre^+^ mice with increasing age (Fig. 2 C-D). Additionally, the isovolumetric relaxation time (IVRT), which is defined as the time between aortic ejection and ventricular filling, was used to evaluate ventricular compliance and relaxation. For the PAD4^fl/fl^ mice, IVRT was significantly increased from 17.30 ± 0.44 ms in the young population to 19.31 ± 0.61 ms in the old group, confirming a decrease in diastolic function with increasing age (Fig. 2 E). In the PAD4^fl/fl^MRP8Cre^+^ mice, IVRT remained constant over the two-year period (young – 17.65 ± 0.78 ms, old – 17.01 ± 0.46 ms) (Fig. 2 E). In the young population, diastolic function, both defined as the E/A ratio and the IVRT, did not differ across the two genotypes of mice.

Vascular function was evaluated in parallel, using PW-doppler echocardiography for measuring systolic pressure in both the aorta and pulmonary artery. No changes were observed in the high pressure system, as AV peak pressure of young PAD4^fl/fl^ (6.68 ± 0.88 mmHg) did not significantly differ from old PAD4^fl/fl^ (7.72 ± 0.72 mmHg) (Fig. 2 F). In line with this, systolic blood pressure in the low-pressure circulation, was not significantly altered in the old population (Fig. 2 G). Clinically, there is an increasing awareness that the right ventricle is of importance with increasing age ^46^. Therefore, right ventricular (RV) function was evaluated based on blood flow in the pulmonary artery. Flow was measured through PW-Doppler and used to measure pulmonary acceleration time (PAT) and pulmonary ejection time (PET). These measurements were used to calculate the ratio of PAT/PET, an established measurement to evaluate RV afterload ^47^. In our mouse model of cardiac aging, no difference was observed in RV function between young and old mice of both genotypes, nor between both genotypes at old age (Fig. 2 H).

### PAD4^fl/fl^MRP8Cre^+^ mice are protected against collagen deposition in the aging heart

Aging is one of the most predisposing factors for fibrotic heart diseases ^48^. Therefore, we assessed cardiac collagen deposition, as a marker for cardiac fibrosis, in young and old PAD4^fl/fl^ and PAD4^fl/fl^MRP8Cre^+^ animals. Corrected heart weight (heart weight divided by tibia length) has previously been used as an overall measure for cardiac hypertrophy development with increasing age ^49^. The corrected heart weight of PAD4^fl/fl^ mice was significantly increased with increasing age (young – 6.92 ± 0.32 mg/mm – n = 12, old – 8.88 ± 0.28 mg/mm – n = 24). The corrected heart weight of PAD4^fl/fl^MRP8Cre^+^ mice did not significantly increase with increasing age (young – 7.73 ± 0.51 mg/mm, n = 9; old – 8.57 ± 0.24 mg/mm – n = 12) (Fig. 3 A). In addition, total cardiac collagen content was assessed and quantified using Masson-Trichrome (Fig. 3 B, D), and Fast Green/ Sirius Red staining (Fig. 3 C, D). As hypothesized based on the decrease in diastolic function in aged PAD4^fl/fl^ mice, these mice showed significantly more cardiac collagen deposition as compared to age-matched PAD4^fl/fl^MRP8Cre^+^ mice (Fig. 3 B, C). In addition, the hearts of young PAD4^fl/fl^ and PAD4^fl/fl^MRP8Cre^+^ mice were analyzed to determine whether the difference in cardiac collagen content in both genotypes was yet present at early developmental stages. At nine weeks of age, collagen content in the heart muscle was comparably low in both genotypes. Therefore, the observed increase in collagen content in the old PAD4^fl/fl^ mice can indeed be classified as an age-related event. Equally interesting, in the old PAD4^fl/fl^MRP8Cre^+^ mice the amount of fibrotic tissue remained similar to that of young PAD4^fl/fl^MRP8Cre^+^ mice for both Masson trichrome staining, as well as the Fast Green/Sirius Red staining, implying that these old mice were protected from age-related changes in myocardial composition and fibrosis.

**Figure 3.**
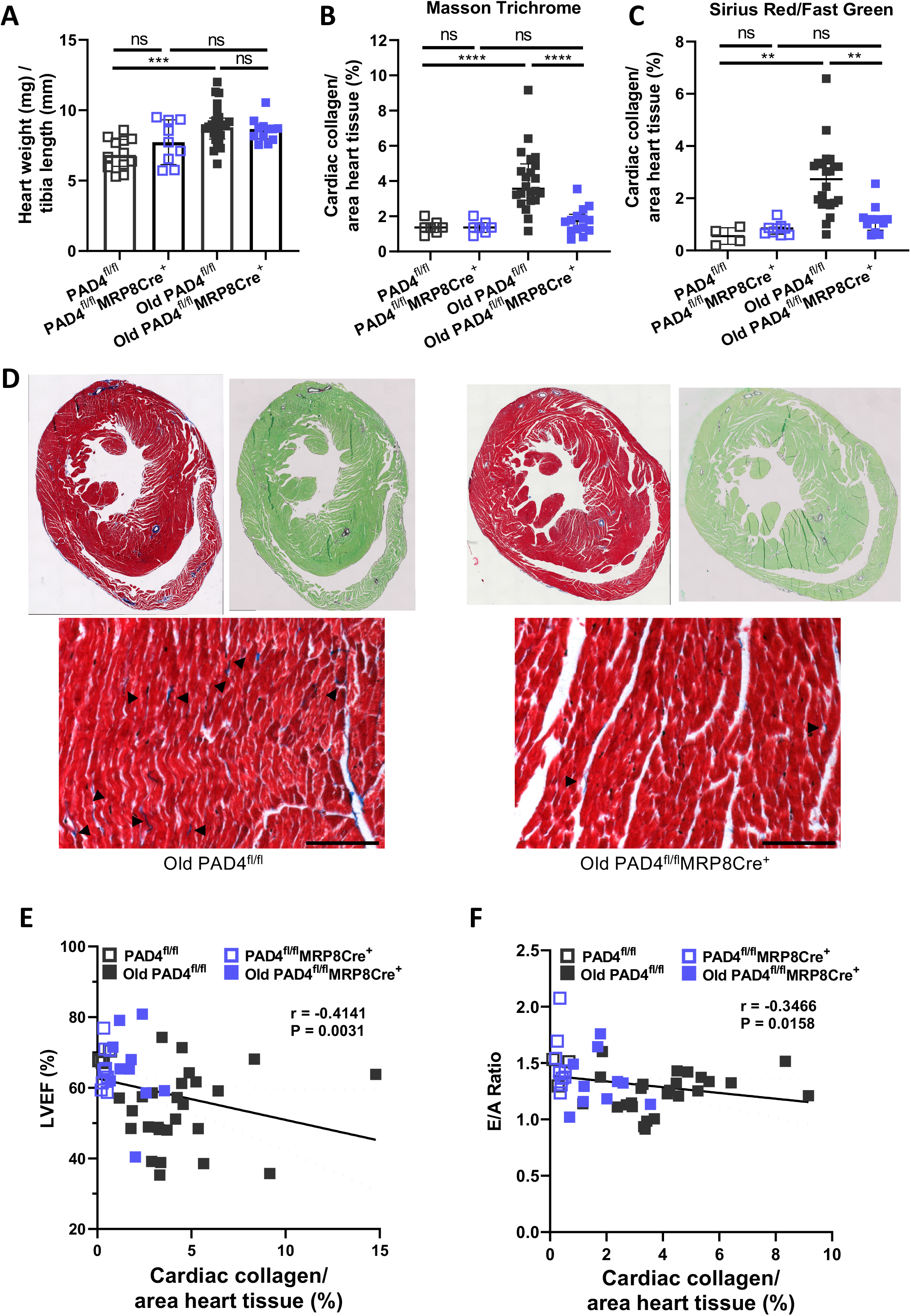
PAD4 deficiency in neutrophils specifically reduces cardiac hypertrophy and collagen deposition with increasing age. (A) Mice were euthanized, perfused and the hearts harvested, and weighed. Simultaneously, tibias were taken and measured. Heart weight of the mice was then corrected for mouse size by dividing by tibia length. Increase in corrected heart weight was taken as a measure for increased cardiac remodeling. (B and C) Cardiac collagen content was assesed by both Masson trichrome and Fast Green/Sirius Red stain in both young and old PAD4^fl/fl^ and PAD4^fl/fl^MRP8Cre^+^ mice. Total collagen was quantified as the percentage of collagen in the whole heart section. (B) The percentage of fibrotic area (blue fibers) in the heart tissue of Masson trichrome stained section was quantified through the color threshold application in ImageJ. Equal color thresholding settings were applied for both young and old mice. (C) Quantification of collagen (Bordeaux fibers) of Fast Green/Sirius Red stained heart tissue of young and old PAD4^fl/fl^ and PAD4^fl/fl^MRP8Cre^+^ mice. Quantification was done through color thresholding on ImageJ. (D) Representative images of Masson trichrome (left) and Fast Green/Sirius Red (right) staining of a horizontal cross-sectional area of the mouse heart of old PAD4^fl/fl^ and PAD4^fl/fl^MRP8Cre^+^ mice. Cardiomyocytes are stained red and green respectively, collagen fibers are stained blue and Bordeaux respectively. Bellow, representative 200X magnifications of Masson Trichrome stained LV wall. Arrowheads indicate the presence of interstitial collagen fibers in the heart tissue. Scale bar: 100 µm. (E and F) Correlation analysis, including all the experimental groups, between the percentage of fibrosis, as determined by the Masson trichrome staining, and systolic cardiac function, given by LVEF (E) or diastolic function, given by E/A ratio (F). NS: not significant, * P < 0.05, ** P < 0.01, *** P < 0.001. Graphs show n = 10 – 32 for young mice, and n = 14 – 25 for old mice.

In order to evaluate whether this observed difference in fibrosis between both genotypes at old age was associated with differences in cardiac function, myocardial fibrosis levels, as determined by Masson trichrome staining, were correlated with both systolic function (via LVEF), and diastolic function (via E/A). Correlation analysis of cardiac fibrosis level with LVEF, showed a significant (P = 0.003) negative correlation with a Spearman correlation coefficient (r) of -0.41 (Fig. 3 E). In addition, correlation analysis of fibrosis level with diastolic function through E/A ratio, revealed a significant (P = 0.016) negative correlation with r = -0.35 (Fig. 3 F). From this we can conclude that neutrophil specific deletion of *Padi4* plays a crucial role in cardiac remodeling induced function deterioration with increasing age.

### Chemotactic gene expression is reduced in old PAD4^fl/fl^MRP8Cre^+^ mice hearts

It is widely known that aging coincides with a number of physiological changes. Gene expression is altered with increasing age, possibly explaining both declines in physical and cognitive abilities ^50^. Since we were mainly interested in cardiac aging, and investigating the complex interplay between neutrophil *Padi4*, age, and cardiac dysfunction, with a particular interest in cardiac fibrosis as an underlying mechanism, we examined gene expression in heart tissue.

We harvested hearts from PAD4^fl/fl^ (n = 13) and *PAD4*^fl/fl^MRP8Cre^+^ (n = 9) mice at 24 mo and isolated mRNA from the heart tissue. This whole mRNA isolate was used to perform quantitative real-time PCR in order to compare expression levels in the aged heart tissue. Expression levels of genes involved in fibrosis (*Agt, Ccl12, Ccn2, Tgfb, Col1a3, Col3a1, Mmp3, Mmp9, Mmp13, Timp1, Timp2, Timp3*, and *Timp4*), inflammation (*Ccl3, Ifng, Il10, Tnf, Smad3*, and *Smad4*), and general HF markers (*Nppa, and Nppb*) were evaluated. This led us to identify equally expressed, upregulated and downregulated genes in old PAD4^fl/fl^MRP8Cre^+^ mice as compared to old PAD4^fl/fl^ mice (Fig. 4). In the hearts of the old PAD4^fl/fl^MRP8Cre^+^ mice, collagen III alpha-1 (*Col3a1*) and tissue inhibitor of metalloprotease 3 (*Timp3*) were upregulated (Fig. 4 – Red), compared to old PAD4^fl/fl^, while C-C motif chemokine 3 (*Ccl3*), C-C motif chemokine 12 (*Ccl12*), CCN family member 2 (*Ccn2*), matrix metalloproteinase 13 (*Mmp13*), and collagen I alpha-3 (*Col1a2*) were downregulated in the old PAD4^fl/fl^MRP8Cre^+^ mice compared to the old PAD4^fl/fl^ mice (Fig. 4 – Blue).

**Figure 4.**
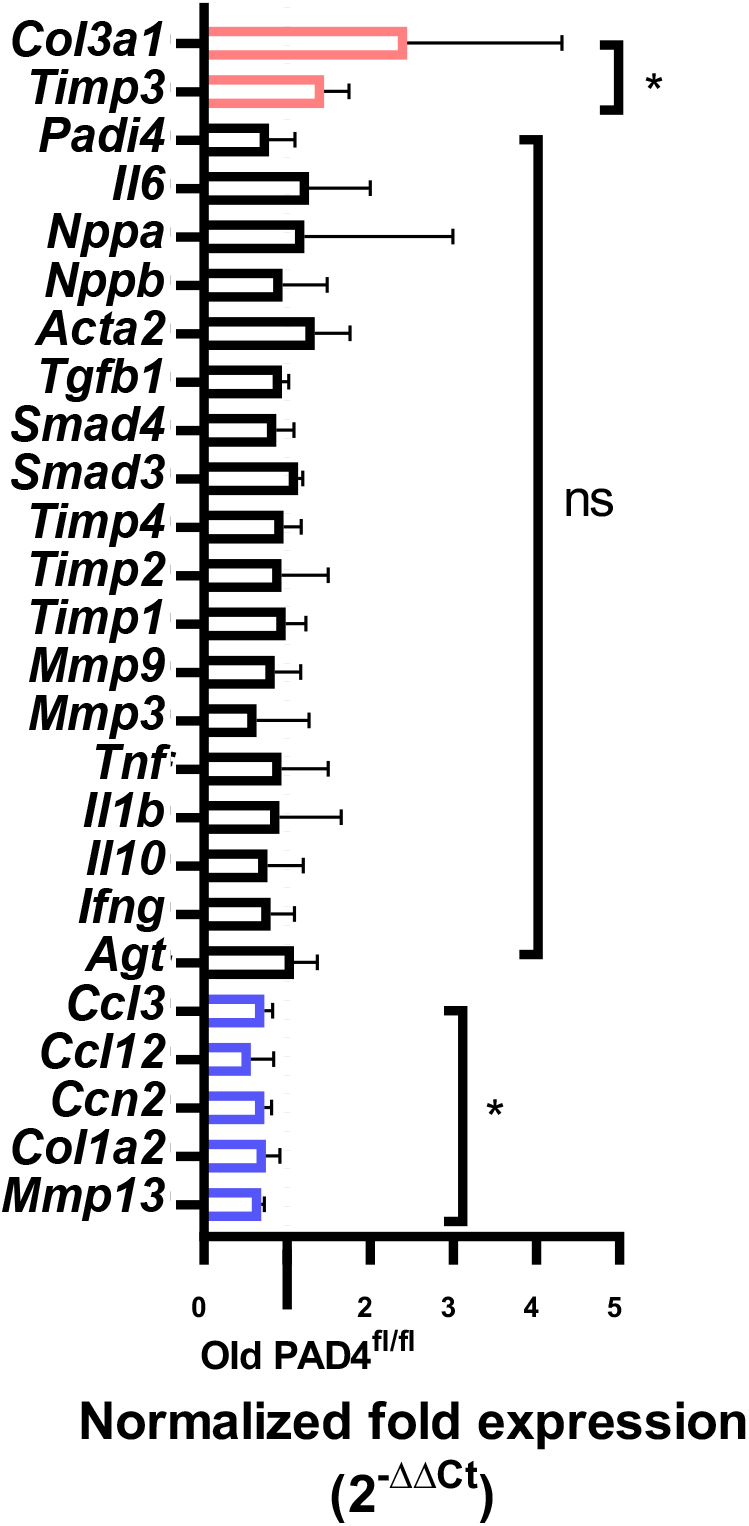
Gene expression in the aging heart is altered due to PAD4 deletion in neutrophils. Quantitative real-time RT-PCR analysis was performed for mRNA expression of several genes in heart tissue from old PAD4^fl/fl^MRP8Cre^+^ mice. Fold expression was calculated with glyceraldehyde 3-phosphate dehydrogenase (GAPDH) as a reference gene, and normalized to mRNA expression in old PAD4^fl/fl^ hearts. Normalized fold expression was taken to be significantly altered (up- or downregulated) when 2^-ΔΔCt^ was significantly different from 1. NS: not significant, * P < 0.05 by Wilcoxon signed rank test with hypothetical value set to one. For old PAD4^fl/fl^ and PAD4^fl/fl^MRP8Cre^+^ groups, n = 8 – 10.

### Plasma levels of chemotactic cytokines are reduced in old PAD4^fl/fl^MRP8Cre^+^ mice

Human physiology is subject to change with increasing age, which will result in a progressive decline in function and increased susceptibility to death ^51^. Apart from decrease in cardiac function, as discussed earlier, one of the most recognized consequences of aging is an alteration in immune function, which can result in a decreased ability to fight infection and constitutive low-grade inflammation (inflammaging) ^52^.

We collected mouse plasma samples of both genotypes at two years of age and compared biomarkers for both aging and inflammation status between groups. As a general marker of aging, cell-free dsDNA (cfDNA) was evaluated ^53^. We could see a clear increase in levels of cfDNA in both old PAD4^fl/fl^ and PAD4^fl/fl^MRP8Cre^+^ mice as compared to their young controls (Fig. 5 A). Previous studies have shown that neutrophils have a higher tendency to go into the process of NETosis, thereby releasing NETs with increasing age ^24^. In order to evaluate neutrophil activation and NETosis, circulating MPO-DNA complexes were evaluated. At 2 years of age, no difference could be observed in plasma levels of these complexes (Old PAD4^fl/fl^ – 4.81 ± 0.24 ng/mL, Old PAD4^fl/fl^MRP8Cre^+^ - 4.40 ± 0.20 ng/mL). Plasma levels of circulating cytokines (IL-15, IL-17A, IL27α, IL-33, CXCL10, CCL2, CCL3, and CXCL2), as well as pro-inflammatory markers (IFN-γ, IL-1β, IL-2, IL-4, IL-5, IL-6, IL-10, IL-12, CXCL1, and TNF-α) were evaluated and compared between genotypes and age groups (Fig. 5 E). Overall, an increase can be seen in circulating cytokines in the old PAD4^fl/fl^ genotype as compared to both young controls and the old PAD4^fl/fl^MRP8Cre^+^ group (Fig. 5 B)). Additionally, we can observe an age-related decrease in some inflammatory cytokines (IFN-γ, IL-10, IL-1β, IL-12, IL-2, and IL-4) in both old PAD4^fl/fl^ and old PAD4^fl/fl^MRP8Cre^+^ mice, while others (IL-6, CXCL1, and TNF-α) demonstrate an age-related increase in plasma levels in the old PAD4^fl/fl^ genotype which is absent in the old PAD4^fl/fl^MRP8Cre^+^ mice (Fig. 5 C). More specifically, plasma levels of interleukin 6 (IL-6) were significantly upregulated in the old PAD4^fl/fl^ as compared to the old PAD4^fl/fl^MRP8Cre^+^ mice (Fig. 5 D). As compared to young control, the plasma levels of tumor necrosis factor α (TNFα) were significantly upregulated in the PAD4^fl/fl^ mice, while this increase was absent in the PAD4^fl/fl^MRP8Cre^+^ genotype (Fig. 5 E). Plasma levels of members of the C-C motif chemokine family were upregulated with increasing age. Both C-C ligand 2 (CCL2) and C-C ligand 3 (CCL3) plasma levels were significantly increased in old mice, as compared to their young controls for both genotypes (Supplemental Fig. 1 A and B). However, members of the C-X-C motif family behaved differently. We could observe a significant increase in the protein concentration of C-X-C ligand 1 (CXCL1) in the plasma of old PAD4^fl/fl^ as compared to young controls, while this increase in plasma levels was absent in the old PAD4^fl/fl^MRP8Cre^+^ genotype (Fig. 5 F). Furthermore, a significant difference could be seen between plasma levels of CXCL1 in both aged groups. Additionally, plasma levels of C-X-C motif ligand 2 (CXCL2) were studied, as this chemokine is important in the recruitment of polymorphonuclear leukocytes, and a known therapeutic target in CVD. Plasma levels of CXCL2 in old PAD4^flf/fl^ mice were not significantly increased compared to young controls (P = 0.07), nor did they differ from old PAD4^fl/fl^MRP8Cre^+^ mice (Supplemental Fig. 1 C).

**Figure 5.**
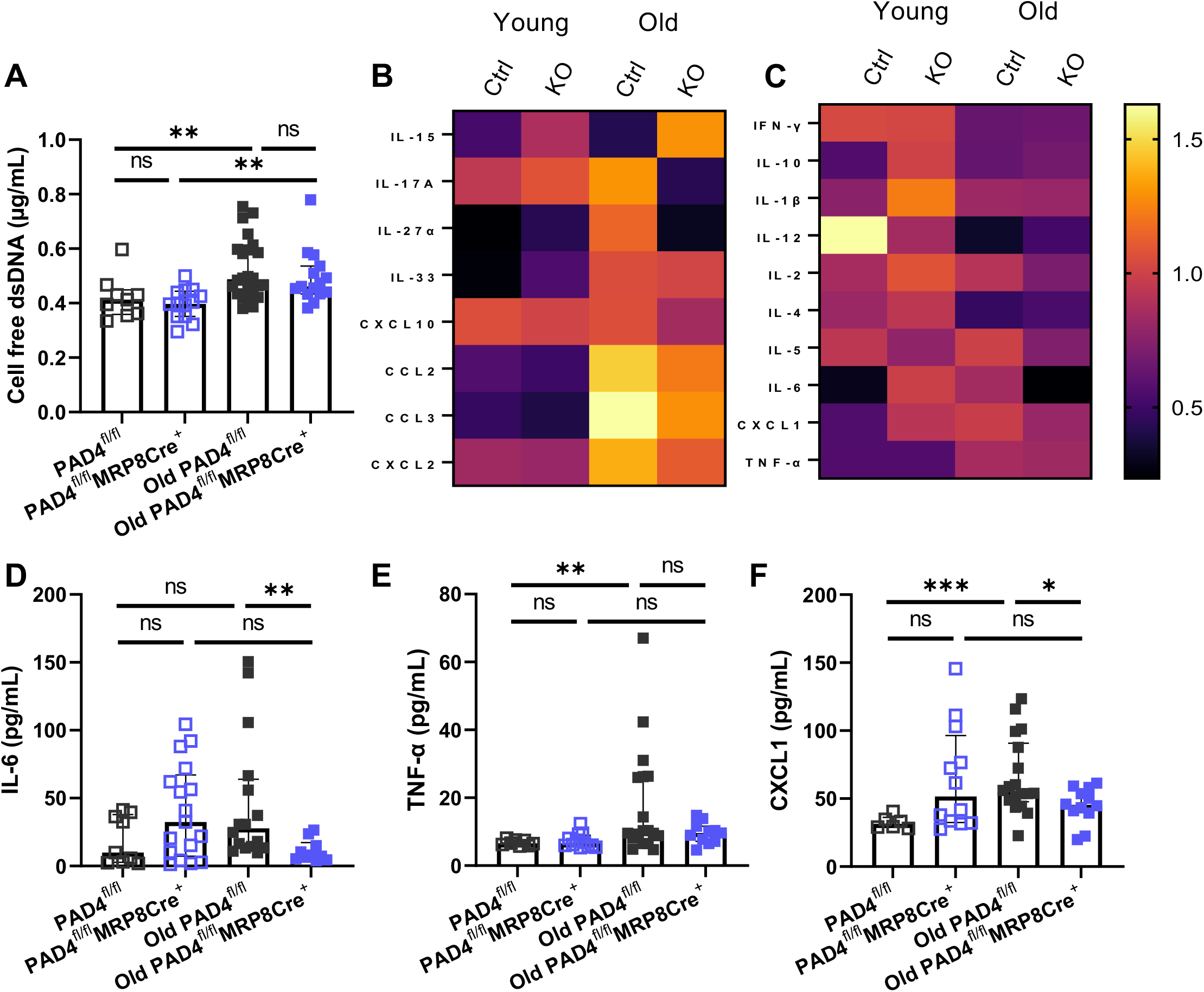
Old PAD4^fl/fl^MRP8Cre^+^ have a decreased pro-inflammatory status with a reduced chemotaxis profile. At day of sacrifice, blood was collected from which plasma was isolated. Plasma biomarkers for aging and inflammation were later determined in batch. (A) Plasma levels of cfDNA. (B) Circulating levels of IL-6. (C) TNF-α concentration as measured in plasma samples. (D) Plasma levels of the neutrophil chemotactic C-X-C Ligand 1 (CXCL1). (E) Heat maps showing relative plasma levels of circulating cytokines and chemokines in the different groups of mice. Rows are corrected by dividing by the average of the row. Black, red, and yellow color indicate increased, equal and decreased plasma levels of the molecule as compared to the average over the two genotypes and two age groups. A: PAD4^fl/fl^, B: PAD4^fl/fl^MRP8Cre^+^, C: old PAD4^fl/fl^, D: old PAD4^fl/fl^MRP8Cre^+^ NS: not significant, * P < 0.05, ** P < 0.01, *** P < 0.001. For young groups n = 10 – 17; for old groups n = 14 – 19.

### Old *PAD4*^fl/fl^MRP8Cre^+^ mice are resistant to metabolically-induced diastolic dysfunction exacerbation

Approximately half of all HF hospital admissions are due to heart failure with preserved ejection fraction (HFpEF) ^54^. The complex clinical pathology that characterizes HFpEF mainly originates from the presence of multiple comorbidities, including obesity, hypertension, and diabetes ^55^. As our aged mice were free from these common complications affecting the human population, we aimed to experimentally recapitulate these in a separate group of animals.

Age-matched groups of two-year-old male mice of both genotypes were started on the ad libitum HFD with 0.5 g/L of L-NAME in the drinking water. After two weeks of HFD and L-NAME exposure, PAD4^fl/fl^ mice experienced a decrease in survival rate, reaching 40% survival after six weeks. On the contrary, PAD4^fl/f^MRP8Cre^+^ mice did not encounter any mortality during the six week period (Fig. 6 A). This resulted in a significant difference in survival between both experimental groups (P = 0.035, PAD4^fl/fl^ n = 9, *PAD4*^fl/fl^MRP8Cre^+^ n = 6). This already suggests a major role of neutrophil PAD4 during environmentally induced HF development, with a specific focus on HFpEF. Systolic and diastolic function were evaluated every 3 weeks, during the 6 week period of increased environmental stress. Systolic function was evaluated based on LVEF measurements, while diastolic function was assessed through E/A ratio. Important to consider is the decreasing number of mice in the PAD4^fl/fl^ group, due to their decreased survival percentage. A clear decrease in LVEF can be observed over the six-week period for the PAD4^fl/fl^ group, reflecting systolic dysfunction. However, this function deterioration differed in the PAD4^fl/fl^MRP8Cre^+^ mice, which showed a cardio-protected phenotype during the first three weeks, followed by a minor decrease in function in the following three weeks. Of note, at week six on the experimental diet, LVEF of PAD4^fl/fl^MRP8Cre^+^ mice was still significantly higher as compared to LVEF of PAD4^fl/fl^ (PAD4^fl/fl^ – 41 ± 4%, PAD4^fl/fl^MRP8Cre^+^ - 59 ± 3%) (Fig. 6 B). For diastolic function, a continued decrease in function can again be observed during the entirety of the exposure period to the metabolic and hypertensive stressors for the old PAD4^fl/fl^ group. After six weeks, E/A ratio reached the threshold value of 1 ± 0.04, showing clear diastolic dysfunction in the PAD4^fl/fl^ mice. For the PAD4^fl/fl^MRP8Cre^+^ genotype, an initially equal decrease in diastolic function can be observed. However, this decline in diastolic function only continued until the third week, after which E/A ratio remained constant (Fig. 6 C). This resulted in a significant difference in diastolic function between both PAD4^fl/fl^ and PAD4^fl/fl^MRP8Cre^+^ mice after six weeks (Fig. 6 C). Collectively, these results indicate an important role of neutrophil PAD4 in the setting of aging accompanied by additional metabolic stress, as is usually the case in human patients with these common risk factors for HF.

**Figure 6.**
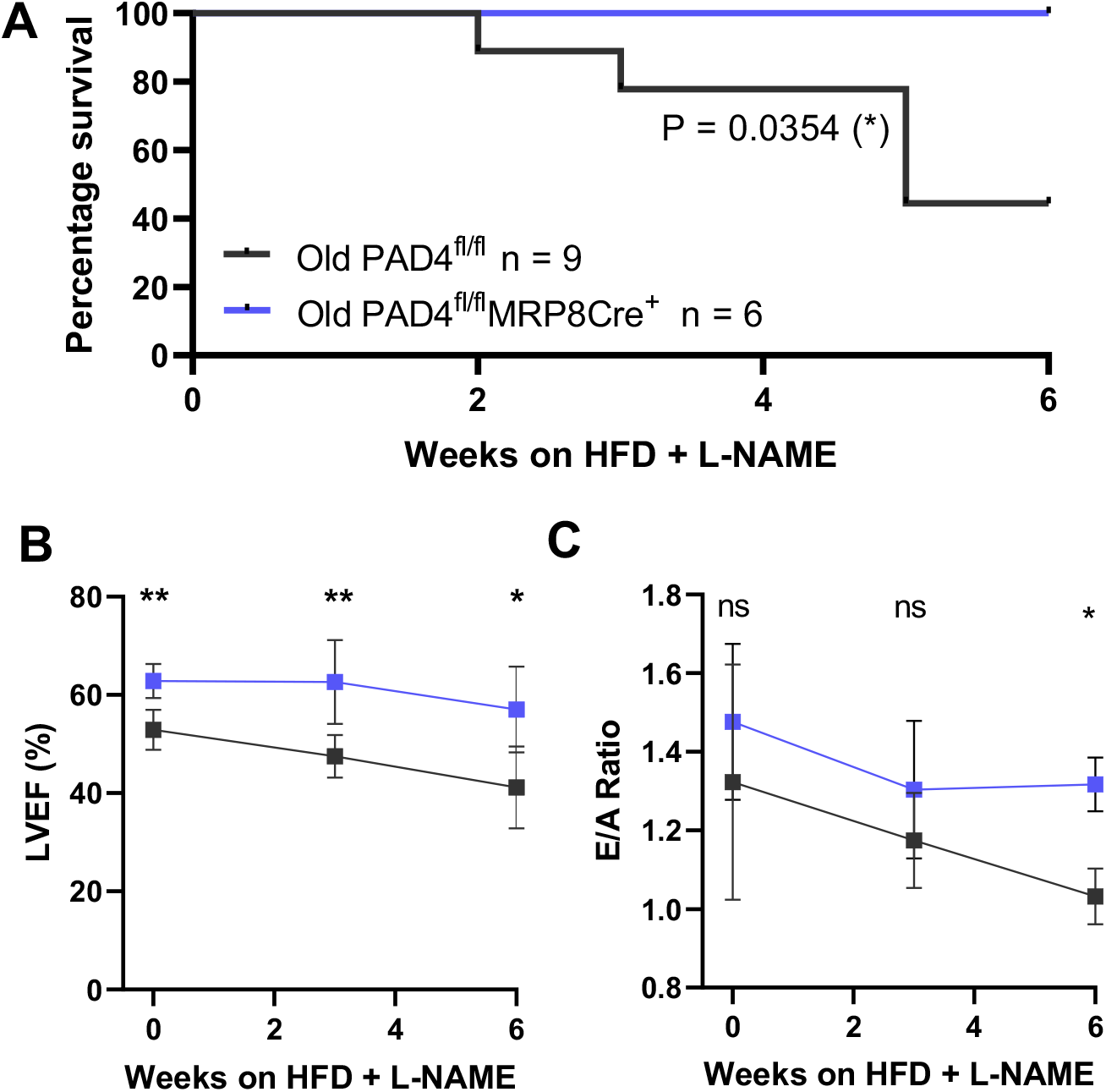
PAD4^fl/fl^MRP8Cre^+^ mice are better protected against metabolic stress at old age. Two-year-old PAD4^fl/fl^ and PAD4^fl/fl^MRP8Cre^+^ mice were exposed to additional metabolic and vascular distress for six weeks by administration of N^ω^-nitro-L-arginine methyl ester (L-NAME) in the drinking water (0.5 g/L), and changing from regular mouse chow to high fat (60% kcal) diet (HFD). (A) Survival curves indicate a decreased probability of survival for old PAD4^fl/fl^ mice when exposed to metabolic and vascular distress. (B and C) Cardiac function evaluation of surviving mice during a 6-week period on which mice were exposed to increased distress. Functional measurements started at day 0, at which mice were 2 years of age. (B) Quantification of systolic function, as evaluated by LVEF, measured from M-mode image taken in the PSAX window. (C) Quantification of diastolic function via E/A ratio, quantified from PW-Doppler images from the A4C view. NS: not significant, * P < 0.05, ** P < 0.01. n = 6 – 9.

## Discussion

Increasing age, specifically being over the age of 65, is the highest risk factor for the development of cardiovascular disease and HF, moreover HF is the leading cause of death in the aged population ^5^. Considering the recent projections of the WHO stating that the population over 60 will increase up to 1.4 billion by 2050, it is clear that HF will remain a major public health burden in the future. In order to ameliorate this, novel strategies for HF prevention and healthy cardiac aging are needed. According to previous research, it is known that systemic deletion of PAD4 resulted in reduced cardiac remodeling and dysfunction at old age.^24^ However, our study revealed the importance of neutrophils, and specifically neutrophil PAD4, as players that interfere with the process of healthy cardiac aging.

Aging is characterized by a gradual reduction in the number of cardiomyocytes, both through necrosis and apoptosis, resulting in dilated cardiomyopathies (DCM).^43^ DCM is considered a serious cardiac disorder which can lead to substantial morbidity and mortality owing to complications such as HF (WHO). In our study, we observed an increase in left ventricular diameter with increasing age in the PAD4^fl/fl^ mice. However, this structural change in the LV was absent in the case of the aged PAD4^fl/fl^MRP8Cre^+^ genotype, with LV internal diameters being comparable to young healthy controls during both peak systole and diastole. In addition, cardiac function of old mice was evaluated for both changes in systolic and diastolic function of the LV, as well as changes in RV afterload. As expected, PAD4^fl/fl^ animals showed an age-dependent decline in heart function comparable to values previously described in wild-type mice.^24^ However, parameters for both systolic function (LVEF), as well as diastolic functional (E/A, and IVRT) in old PAD4^fl/fl^MRP8Cre^+^ mice did not show signs of cardiac dysfunction. This decrease in function deterioration in PAD4^fl/fl^MRP8Cre^+^ mice is consistent with the previously established phenotype in systemic PAD4^-/-^ mice,^24^ indicating that the phenotype is neutrophil-driven. The absence of systolic dysfunction in the old PAD4^fl/fl^MRP8Cre^+^ genotype can be tied to the absence in LV dilation in these mice, highlighting that neutrophil specific deletion of *Padi4* protects against age induced LV dilation and LV systolic dysfunction. LV dilation during aging can be explained by both apoptosis and necrosis of the cardiomyocytes.^39-42^ It has been shown that PAD4-deficiency results in smaller infarcts in ischemia/reperfusion (I/R) injury studies ^29^, and that NET infiltration in the myocardium during acute cardiac injury results in apoptosis of cardiomyocytes ^56^. Therefore, neutrophils, and NETs may promote similar processes of cardiomyocyte death during chronic inflammation, as is the case during natural aging.

Previous experimental as well as clinical studies have already provided extensive evidence suggesting that the aging heart undergoes aberrant fibrotic remodeling ^57^. During this process collagen content in the heart increases, leading to progressive stiffening of the ventricles and impaired diastolic function. Cardiac collagen content was assessed as a measure for cardiac fibrosis in both young and old mice. We found an age-dependent increase in cardiac collagen content in our PAD4^fl/fl^ genotype, which is consistent with what was previously described in other reports ^24, 57^. Strikingly, in our setting of neutrophil specific deletion of *Padi4* we could observe a cardiac collagen content which did not differ between young and old groups. This protection from age induced cardiac collagen deposition was also observed in the aged systemic PAD4 deletion mice, previously described ^24^. During the process of fibrotic remodeling, the trans-differentiation towards, and migration of myofibroblasts at/to the site of inflammation is of major importance ^8^. Activation of fibroblasts towards α-SMA producing myofibroblasts has been shown to be influenced by NETs in vitro ^58^. This activation of fibroblasts in culture resulted in increased connective tissue growth factor production, collagen deposition, and fibroblast migration ^58^. NETs may be able to promote similar processes in the myocardium during the process of natural aging, thereby promoting cardiac fibrosis and further remodeling.

Next, because of the protection against age-induced cardiac collagen deposition in the PAD4^fl/fl^MRP8Cre^+^ mice, we studied the downstream pathways impacted by neutrophil PAD4, and how they could be involved in collagen deposition in the heart. Fibrotic remodeling is known to be a complex multi-stage process, in which bone marrow-derived leukocytes are crucial ^10^. When looking at changes in cardiac gene expression between hearts of both genotypes at old age, changes in gene expression could be observed as a result of neutrophil *Padi4* deletion. The simultaneous upregulation of *Col3a1* and downregulation of *Col1a2* seem contradictory; however, these two types of collagen have distinct molecular makeup and functional properties. Type I collagen fibrils are the most abundant, and are considered stiff structures, rendering tissue rigid and durable ^59, 60^. On the other hand, type III collagen fibrils are more thin than type I, and are present in high concentrations in tissues which require elastic properties ^61^. *Timp3*, upregulated in PAD4^fl/fl^MRP8Cre^+^ mice, has been shown to be both involved in cardiac fibrosis clearance, as well as amelioration of inflammation, with TIMP3^-/-^ mice showing both increased cardiac collagen content, and a significant increase of cardiac infiltrated neutrophils in a model of angiotensin II induced cardiac hypertrophy ^62^. Additionally, high expression levels of *Timp3* result in protection of cardiomyocytes against apoptosis in acute injury settings ^63^, while TIMP*3* deficiency leads to dilated cardiomyopathy in mice ^64^. On the other hand, *Mmp13* is downregulated in the hearts of PAD4^fl/fl^MRP8Cre^+^ mice as compared to PAD4^fl/fl^. MMP13 is involved in ECM degradation, including triple helical collagens containing type I, II, and III, and is shown to be upregulated in hypertensive hearts of rats ^65^, and is targeted by doxycycline administration in human patients suffering from HF ^66^. In accordance, it was shown that inhibition of MMP13 in a model of LV pressure overload in mice reduced cardiac hypertrophy and resulted in protection against hypertension induced cardiac dysfunction, suggesting that MMP13 plays a detrimental role in pressure overload-induced HF ^67^.

With respect to inflammation-related gene expression, we could identify several inflammatory markers which were downregulated in old PAD4^fl/fl^MRP8Cre^+^ mice as compared to the old PAD4^fl/fl^. We saw a downregulation of *Ccl3* in the old PAD4^fl/fl^MRP8Cre^+^ mice. This C-C motif chemokine is known to be upregulated in HF patients ^68^, and causes intracellular calcium release and recruitment of neutrophils ^69^, thus having a possible role in NET formation and cardiac inflammation, respectively. Interestingly, *Ccl12* was downregulated in the neutrophil PAD4 knockout group. This chemotactic factor attracts eosinophils, monocytes, and lymphocytes to sites of inflammation. In addition, it has been shown that this chemokine is responsible for fibrocyte recruitment towards the lung tissue under acute injury settings, orchestrating a fibrotic response at the site of injury ^70^. Finally, we discovered a potential interaction between neutrophil *Padi4* and *Ccn2*. This CCN family member 2 promotes fibrosis development, and is involved in the aging process. From recent research, it was elucidated that CCN2 is an autocrine regulator of fibroblast activation, modulating fibrosis development in the heart ^71^. Interestingly, we found that in our old PAD4^fl/fl^MRP8Cre^+^ mice, this gene was downregulated. All these changes in gene expression between old PAD4^fl/fl^MRP8Cre^+^ mice and PAD4^fl/fl^ mice corroborate both the protection against cardiac remodeling and the conservation of cardiac function in old *PAD4*^fl/fl^MRP8Cre^+^ mice, as compared to old PAD4^fl/fl^ mice.

Aging is associated with an elevated inflammatory activity, a phenomenon previously described as inflammaging. This is associated with increased inflammatory activity in the blood, including elevated levels of circulating TNF-α ^72^ and IL-6 ^73^. In concordance with these previously described results, we found an elevated level of TNF-α in the plasma of old PAD4^fl/fl^ mice, as compared to young controls. Interestingly, this age-dependent increase was absent in the PAD4^fl/fl^MRP8Cre^+^ genotype. TNF-α has been shown to induce neutrophils of healthy subject towards NET release through the activation of NADPH oxidase and MPO ^74^. Additionally, a difference could be seen between plasma levels of IL-6 when comparing both genotypes at old age, with PAD4^fl/fl^ mice showing increased levels as compared to PAD4^fl/fl^MRP8Cre^+^ mice. With advancing age, plasma levels of IL-6 become detectable in the absence of infection ^73^. Additionally, IL-6 causes compromised tissue repair by shifting acute inflammation into a more profibrotic state trough the induction of Th1 cell responses due to recurrent inflammation ^75^. Apart from their clear roles in inflammation and fibrosis, both TNF-α and IL-6 have been described as mediators of HF progression. TNF-α has been shown to be a mediator of myocardial dysfunction leading to HF progression ^76^, while IL-6 was elevated in a cohort study of HF patients, were it was associated with reduced LVEF and poorer clinical outcomes ^77^.

In addition to increases in inflammation status in old PAD4^fl/fl^ mice, our study revealed that deletion of neutrophil *Padi4* prevents the age-induced increase in plasma CXCL1 levels. Ligand 1 of the C-X-C motif chemokines is also known as neutrophil activating protein 3, and the homologue of IL-8 in humans, and is mainly produced by neutrophils, macrophages and fibroblasts ^78^. CXCL1 is a pro-inflammatory factor and is mainly involved in neutrophil chemotaxis ^79^. Moreover, the CXCL1-CXCR2 axis is involved in neutrophil degranulation and NET release ^79, 80^. Recent investigations demonstrated that disruption of this signaling axis has the possibility to attenuate cardiac fibrosis, hypertrophy, and dysfunction in hypertensive rat hearts ^81^. Altogether, neutrophil PAD4 can be proposed to play a role in neutrophil-dependent leukocyte recruitment, and fibrosis in the heart muscle at old age. Finally, deletion of neutrophil *Padi4* played a significant role in survival during a period of increased metabolic and mechanical stress. Geriatric obesity has long been recognized as a health issue in developed countries, and is now rising in developing countries as well. During metabolic stress, as is the case during high caloric intake, neutrophils have been shown to be increased in absolute numbers, as well as being activated in circulation ^82, 83^. The ability to form NETs is disrupted upon deletion of neutrophil *Padi4*, therefore inhibiting a possible pathway by which neutrophils can cause cardiac damage in setting of obesity.

In summary, understanding the mechanisms involved in age-induced organ remodeling and dysfunction are essential for developing proper treatment and care for the elderly population. Our study has showed that neutrophils, and more specifically neutrophil PAD4, are crucial during the process of unhealthy cardiac aging. This makes neutrophil PAD4 a valid therapeutic target for both heart failure prevention, and progression treatments during settings of chronic inflammation, as is the case during natural aging.

## Conclusion

The absence of neutrophil peptidylarginine deiminase 4 (*Padi4*) during the process of natural aging, results in healthy cardiac function at old age in mice. The absence of function deterioration can be explained by reduced cardiac fibrosis, which can be justified by a possible reduction in recruitment of inflammatory cells to the myocardium, and a reduction in overall inflammation status due to the absence of neutrophil *Padi4*. Further investigation in this process of immune cell recruitment towards the heart muscle might reveal novel therapeutic strategies which can prevent the irreversible scarring of the heart muscle, a process crucial in heart failure development and progression.

## Authors’ contributions

Conceptualization: S.V.B. and K.M.; Methodology: S.V.B., S.K., J.V.W., and P.C.; Analysis: S.V.B., S.K., and K.M.; Investigation: S.V.B., resources: T.W., and K.M.; writing – original draft: S.V.B.; Writing – review and editing: S.V.B., S.K., J.V.W., P.C., L.F., T.W., and K.M.; Supervision: K.M., project administration: S.V.B., and K.M., funding acquisition: T.V. and K.M.

## Acknowledgements

We would like to thank the members of the Martinod group in the Center for Molecular and Vascular Biology, Department of Cardiovascular Science, KU Leuven, Belgium for helpful discussions and training in experimental techniques.

