## Supplemental files for "Neutrophil Peptidylarginine Deiminase 4 is Essential for Detrimental Age-related Cardiac Remodeling & Dysfunction in Mice"

### Slide 1
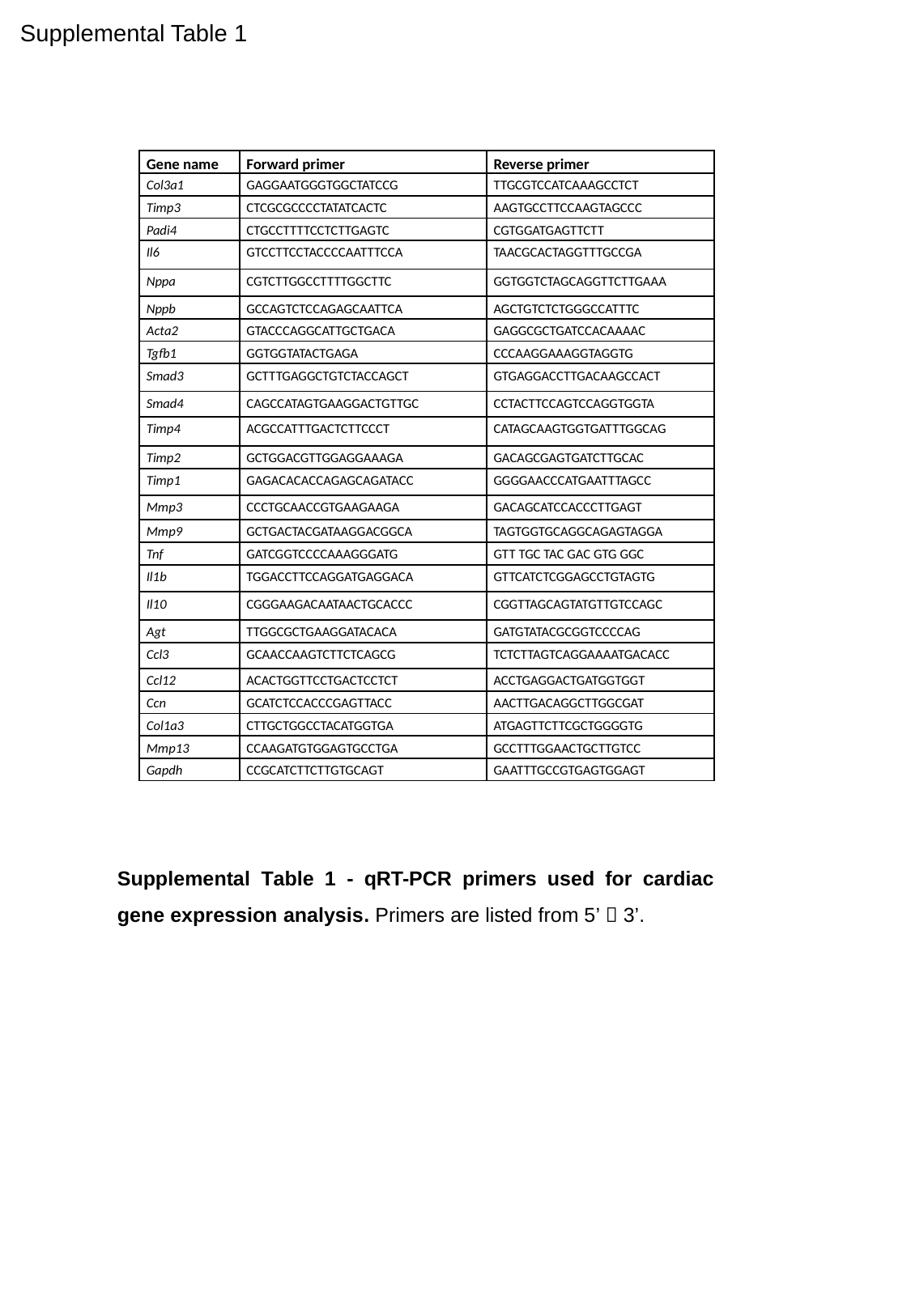

Supplemental Table 1
| Gene name | Forward primer | Reverse primer |
| --- | --- | --- |
| Col3a1 | GAGGAATGGGTGGCTATCCG | TTGCGTCCATCAAAGCCTCT |
| Timp3 | CTCGCGCCCCTATATCACTC | AAGTGCCTTCCAAGTAGCCC |
| Padi4 | CTGCCTTTTCCTCTTGAGTC | CGTGGATGAGTTCTT |
| Il6 | GTCCTTCCTACCCCAATTTCCA | TAACGCACTAGGTTTGCCGA |
| Nppa | CGTCTTGGCCTTTTGGCTTC | GGTGGTCTAGCAGGTTCTTGAAA |
| Nppb | GCCAGTCTCCAGAGCAATTCA | AGCTGTCTCTGGGCCATTTC |
| Acta2 | GTACCCAGGCATTGCTGACA | GAGGCGCTGATCCACAAAAC |
| Tgfb1 | GGTGGTATACTGAGA | CCCAAGGAAAGGTAGGTG |
| Smad3 | GCTTTGAGGCTGTCTACCAGCT | GTGAGGACCTTGACAAGCCACT |
| Smad4 | CAGCCATAGTGAAGGACTGTTGC | CCTACTTCCAGTCCAGGTGGTA |
| Timp4 | ACGCCATTTGACTCTTCCCT | CATAGCAAGTGGTGATTTGGCAG |
| Timp2 | GCTGGACGTTGGAGGAAAGA | GACAGCGAGTGATCTTGCAC |
| Timp1 | GAGACACACCAGAGCAGATACC | GGGGAACCCATGAATTTAGCC |
| Mmp3 | CCCTGCAACCGTGAAGAAGA | GACAGCATCCACCCTTGAGT |
| Mmp9 | GCTGACTACGATAAGGACGGCA | TAGTGGTGCAGGCAGAGTAGGA |
| Tnf | GATCGGTCCCCAAAGGGATG | GTT TGC TAC GAC GTG GGC |
| Il1b | TGGACCTTCCAGGATGAGGACA | GTTCATCTCGGAGCCTGTAGTG |
| Il10 | CGGGAAGACAATAACTGCACCC | CGGTTAGCAGTATGTTGTCCAGC |
| Agt | TTGGCGCTGAAGGATACACA | GATGTATACGCGGTCCCCAG |
| Ccl3 | GCAACCAAGTCTTCTCAGCG | TCTCTTAGTCAGGAAAATGACACC |
| Ccl12 | ACACTGGTTCCTGACTCCTCT | ACCTGAGGACTGATGGTGGT |
| Ccn | GCATCTCCACCCGAGTTACC | AACTTGACAGGCTTGGCGAT |
| Col1a3 | CTTGCTGGCCTACATGGTGA | ATGAGTTCTTCGCTGGGGTG |
| Mmp13 | CCAAGATGTGGAGTGCCTGA | GCCTTTGGAACTGCTTGTCC |
| Gapdh | CCGCATCTTCTTGTGCAGT | GAATTTGCCGTGAGTGGAGT |
Supplemental Table 1 - qRT-PCR primers used for cardiac gene expression analysis. Primers are listed from 5’ 🡪 3’.

### Slide 2
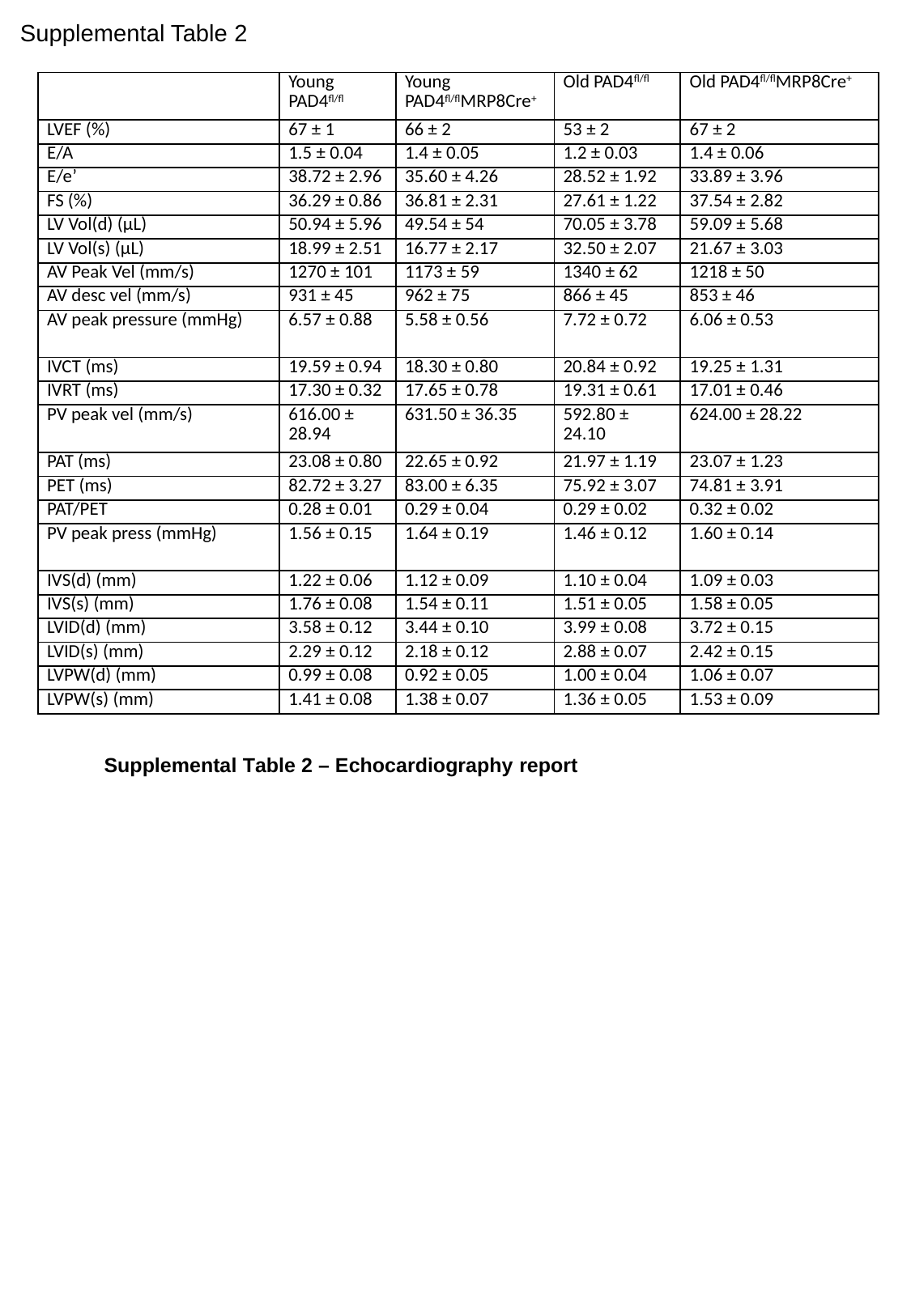

Supplemental Table 2
| | Young PAD4^fl/fl^ | Young PAD4^fl/fl^MRP8Cre^+^ | Old PAD4^fl/fl^ | Old PAD4^fl/fl^MRP8Cre^+^ |
| --- | --- | --- | --- | --- |
| LVEF (%) | 67 ± 1 | 66 ± 2 | 53 ± 2 | 67 ± 2 |
| E/A | 1.5 ± 0.04 | 1.4 ± 0.05 | 1.2 ± 0.03 | 1.4 ± 0.06 |
| E/e’ | 38.72 ± 2.96 | 35.60 ± 4.26 | 28.52 ± 1.92 | 33.89 ± 3.96 |
| FS (%) | 36.29 ± 0.86 | 36.81 ± 2.31 | 27.61 ± 1.22 | 37.54 ± 2.82 |
| LV Vol(d) (µL) | 50.94 ± 5.96 | 49.54 ± 54 | 70.05 ± 3.78 | 59.09 ± 5.68 |
| LV Vol(s) (µL) | 18.99 ± 2.51 | 16.77 ± 2.17 | 32.50 ± 2.07 | 21.67 ± 3.03 |
| AV Peak Vel (mm/s) | 1270 ± 101 | 1173 ± 59 | 1340 ± 62 | 1218 ± 50 |
| AV desc vel (mm/s) | 931 ± 45 | 962 ± 75 | 866 ± 45 | 853 ± 46 |
| AV peak pressure (mmHg) | 6.57 ± 0.88 | 5.58 ± 0.56 | 7.72 ± 0.72 | 6.06 ± 0.53 |
| IVCT (ms) | 19.59 ± 0.94 | 18.30 ± 0.80 | 20.84 ± 0.92 | 19.25 ± 1.31 |
| IVRT (ms) | 17.30 ± 0.32 | 17.65 ± 0.78 | 19.31 ± 0.61 | 17.01 ± 0.46 |
| PV peak vel (mm/s) | 616.00 ± 28.94 | 631.50 ± 36.35 | 592.80 ± 24.10 | 624.00 ± 28.22 |
| PAT (ms) | 23.08 ± 0.80 | 22.65 ± 0.92 | 21.97 ± 1.19 | 23.07 ± 1.23 |
| PET (ms) | 82.72 ± 3.27 | 83.00 ± 6.35 | 75.92 ± 3.07 | 74.81 ± 3.91 |
| PAT/PET | 0.28 ± 0.01 | 0.29 ± 0.04 | 0.29 ± 0.02 | 0.32 ± 0.02 |
| PV peak press (mmHg) | 1.56 ± 0.15 | 1.64 ± 0.19 | 1.46 ± 0.12 | 1.60 ± 0.14 |
| IVS(d) (mm) | 1.22 ± 0.06 | 1.12 ± 0.09 | 1.10 ± 0.04 | 1.09 ± 0.03 |
| IVS(s) (mm) | 1.76 ± 0.08 | 1.54 ± 0.11 | 1.51 ± 0.05 | 1.58 ± 0.05 |
| LVID(d) (mm) | 3.58 ± 0.12 | 3.44 ± 0.10 | 3.99 ± 0.08 | 3.72 ± 0.15 |
| LVID(s) (mm) | 2.29 ± 0.12 | 2.18 ± 0.12 | 2.88 ± 0.07 | 2.42 ± 0.15 |
| LVPW(d) (mm) | 0.99 ± 0.08 | 0.92 ± 0.05 | 1.00 ± 0.04 | 1.06 ± 0.07 |
| LVPW(s) (mm) | 1.41 ± 0.08 | 1.38 ± 0.07 | 1.36 ± 0.05 | 1.53 ± 0.09 |
Supplemental Table 2 – Echocardiography report

### Slide 3
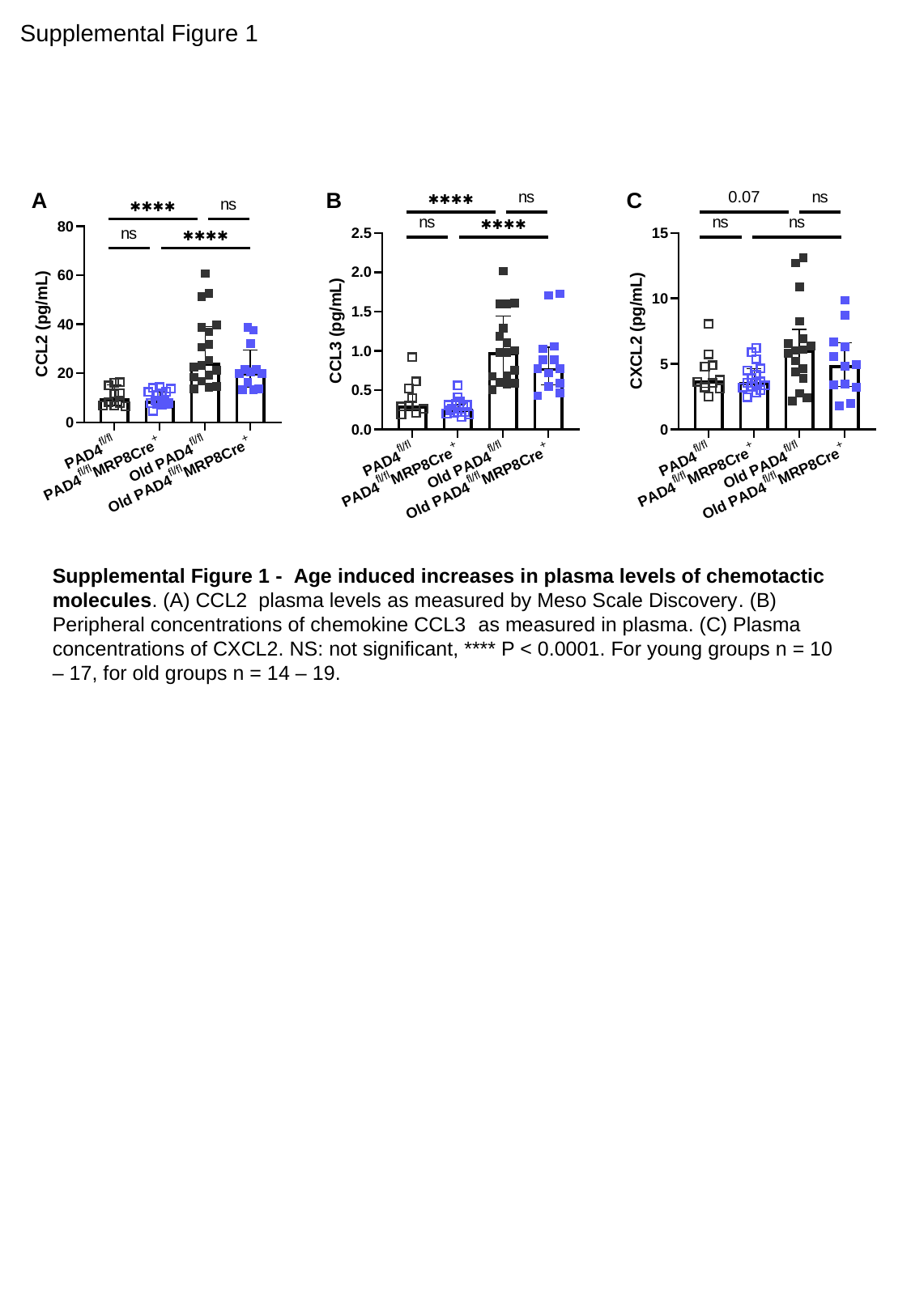

Supplemental Figure 1
A
B
C
Supplemental Figure 1 - Age induced increases in plasma levels of chemotactic molecules. (A) CCL2  plasma levels as measured by Meso Scale Discovery. (B) Peripheral concentrations of chemokine CCL3  as measured in plasma. (C) Plasma concentrations of CXCL2. NS: not significant, **** P < 0.0001. For young groups n = 10 – 17, for old groups n = 14 – 19.
